# Quantitative Analysis of Drug Efficacy on *C. elegans* Models for Neuromuscular Diseases

**DOI:** 10.1101/2021.01.21.427562

**Authors:** Samuel Sofela, Sarah Sahloul, Yong-Ak Song

## Abstract

*Caenorhabditis elegans* has emerged as a powerful model organism for drug screening due to its cellular simplicity, genetic amenability and homology to humans combined with its small size and low cost. Currently, high-throughput drug screening assays are mostly based on image-based phenotyping not exploiting key locomotory parameters of this multicellular model with muscles such as its thrashing force, a critical parameter when screening drugs for muscle-related diseases. In this study, we demonstrated the use of a micropillar-based force assay chip in combination with an imaging assay to evaluate the efficacy of various drugs currently used in treatment of neuromuscular diseases. Using this two-dimensional approach, we showed that the force assay was generally more sensitive in measuring efficacy of drug treatment in Duchenne Muscular Dystrophy and Parkinson’s Disease mutant worms as well as partly in Amyotrophic Lateral Sclerosis model. These results underline the potential of our force assay chip in screening of potential drug candidates for the treatment of neuromuscular diseases when combined with an imaging assay in a two-dimensional analysis approach.

## Introduction

Neuromuscular diseases consist of a diverse group of medical conditions that mainly affect the one or more parts of the neuromuscular unit such as skeletal muscle, motor neurons, peripheral nerves and neuromuscular synapses[1]. These diseases which can be hereditary or acquired, affect as many as 1 in 3,000 people[2] and can be attributed to neurodegenerative diseases like Parkinson’s disease (PD)[3] and Duchenne muscular dystrophies (DMD)[4]. A common effect of most neuromuscular disorders is locomotion due to muscle wasting, weakness and disuse[5].

*Caenorhabditis elegans* has been utilized as a relevant disease model to explain the intricacies of cellular processes and in the search for drugs for treatment of neuromuscular diseases[6]. In modeling of neuromuscular diseases, it is important that neuronal cellular functions and muscle structure are well conserved if drugs are to be translated to humans. As such, several drugs used in to treat neuromuscular and neurodegenerative diseases have proven effective in phenotyping *C. elegans* for validation[7–9].

Of particular interest is the locomotory mechanism of the *C. elegans*, which is involved in most of the worm’s behavior, and has been used in genetic analysis to score phenotypes linked to neuromodulatory and structural defects[10]. The locomotory dynamics in *C. elegans* is not only derived from a combination of neuromuscular control systems but also from the coordination of internal control and physical properties of the worm[11,12]. Its locomotory behavior has mostly been quantified through measuring locomotive parameters, such as worm velocity, thrashing frequency, and number of bends[13]. While these parameters have proven beneficial, they are indirect phenotypes and provide a different aspect of worm physiology compared to force assays[14].

Due to the similar length scale, microfluidics technology has emerged as a powerful platform in the phenotyping of *C. elegans*[15–17]. Cornaglia et. al reported a multi-functional microfluidic platform for automated worm culture, immobilization and long-term imaging of Amyotrophic Lateral Sclerosis (ALS) and Huntington disease mutant worms[18]. In another study[19], long-term swimming exercise was used as a behavioral phenotype to understand the impact of exercise on locomotory performance in Alzheimer’s and Huntington’s disease model animals. Salam et al[20] utilized electrotaxis response of *C. elegans* to analyze the worm’s response to neurotoxic and neuroprotective compounds in modeling PD. The aforementioned assays have contributed to better understanding and assessment of drug treatments for neuromuscular and neurodegenerative diseases. However, there was no direct characterization of muscle strength as the ability to generate the maximal amount of muscle force which can be correlated with clinical studies in humans. To this end, there have been several studies that quantified the force exerted by body wall muscles using elastomeric micropillars[21–24], however, only Hewitt et al used a neuromuscular disease model[14]. In their study, they reported the use of their micropillar-based force measurement system, called Nemaflex, to study the muscle strength of DMD mutant worms before and after drug treatment. Using their device with free-moving worms, they were able to show that adult *dys-1(eg33)* mutants were weaker than wild-type worms while *dys-1(cx18)* mutants exhibited similar muscle force as wild-type worms. While these studies successfully quantified the muscle force, the use of free-moving worms could introduce complexities for tracking the worm which could pose major challenges for multiplexing. Although tracking of free-moving worms in micropillar array has been demonstrated[25], its throughput is still limited.

In this study, we performed a two-dimensional analysis of drug efficacy using *C. elegans* as neuromuscular disease models. For the first-dimensional analysis, we utilized a microfluidic chip with an integrated array of elastomeric micropillars to partially immobilize a *C. elegans* and quantitatively measure its thrashing force before and after drug treatment. This thrashing force assay device in polydimethylsiloxane (PDMS) allowed us to quantify the force exerted by partially immobilized worms in various developmental stages independent of their trapping orientation[24]. To evaluate the muscle strength degradation in neuromuscular disease (NMD) model worms, we used three mutant worms: DMD (LS587), ALS (AM725), and Parkinson’s Disease (NL5901). Using these models in the force assay device, we examined the efficacy of six representative drugs for the treatment of neuromuscular diseases. In the second-dimensional analysis, we performed a quantitative image analysis of the protein aggregation and morphological studies of the body wall muscles before and after drug treatment on an agarose pad following standard protocol and validated the force measurement data. In this way, we could quantify the efficacy of the drugs on the muscle force and corroborate the force data with morphological studies of the protein aggregation and actin filament structures in the body wall muscles for validation. With its simple and scalable design, our force assay chip has facilitated a highly quantitative and sensitive biophysical phenotyping of *C. elegans* without biases to assess efficacy of various drugs on muscle strength.

## Materials and Methods

### Worm Strain

Three transgenic strains were used in this study to model neuromuscular diseases: LS587 *(dys-1(cx18) I; hlh-1(cc561) II)* as DMD model, AM725 *(rmIs290[unc-54p::Hsa-sod-1(127X)::YFP]*) as ALS model, and NL5901 *([unc54p::alphasynuclein: :YFP + unc-119(+)])* as PD model. All strains were obtained Caenorhabditis Genetics Center (University of Minnesota, Minnesota, MN). LS587 strain was maintained at 15°C, while AM725 and NL5901 were maintained at 20 °C. Worms were grown using the standard nematode growth medium seeded with *Escherichia coli* [26]. Sodium hypochlorite treatment was done to obtain embryos from gravid adult animals, eggs were allowed to hatch at room temperature except for LS587 strain which was kept at 15°C.

### Drug Treatment and Culture

For each strain, two different drugs were evaluated. For the DMD model, LS587, two common drugs were tested: prednisone and melatonin. Each drug was added to the Nematode Growth Media (NGM) with a final concentration of 0.37mM prednisone diluted in 0.062% DMSO and 1 mM melatonin. Both drug concentrations were selected based on previous drug treatment study on *C. elegans* [14]. Worms grown on NGM mixed with the required drug starting from L1 stage for ∼3 days and 17 hours at 15°C until L4 stage, and for 4 days and ∼17 hours at 15°C until young adult stage. Control plates for prednisone contained 0.062% DMSO.

ALS model worms were treated with doxycycline and riluzole at two different concentrations. Doxycycline was added to the NGM plates to obtain a final concentration of 10.5 μM and 32 μM. Worms were cultured at 20 °C. The control group was kept for ∼2 days and 4 hours to reach young adult stage. The worms treated with 10.5 μM doxycycline required additional ∼5 hours and those treated with 32 μM doxycycline additional ∼9 hours compared to the control group. This delay in growth is known as a side effect by doxycycline[18]. The higher dosage of doxycycline (32 μM) was tested on SJ4100 and showed a decrease in the average size of aggregates[18]. For comparison, we also selected a lower dosage of the drug as well. Worms treated with riluzole were maintained at 20 °C until L4 stage and then moved to liquid culture, the S basal medium containing the drug, until reaching young adult stage. Both the drug treated group and the control group contained 0.5% DMSO which was used as a solvent for riluzole. The control sample was incubated for ∼1 day and 21 hours while for drug treated ones at 30 μM and 100 μM were incubated by ∼2 hours longer before it was shifted to liquid culture for drug treatment. In liquid culture, both the control and 30 μM riluzole treated worms were incubated for ∼1 day with shaking at 100 rpm in 20 °C in 96 well plate, while 100 μM riluzole treated ones required additional incubation by ∼3 hours to reach young adult stage[13]. Riluzole treatment was shifted to liquid media in S basal due to the low solubility of riluzol in NGM similar to the study by Ikenaka et al.[13].

Levodopa and pramipexole were used to treat NL5901. Levodopa was mixed with NGM plates for the final concentrations of 0.7 mM and 2 mM with 0.5% DMSO[27]. The control group also was treated with 0.5% DMSO. Worms were maintained at 20 °C from L1 stage until young adult for ∼2 days and 5 hours. Pramipexole was prepared in M9 buffer and spread on top of the NGM plates with a final concentration of 2.5 mM and 5 mM. 5mM pramipexole has been reported in the treatment of C. elegans[28], and the lower dosage of 2.5 mM was used because excess amounts of pramipexole may result in over stimulation of *C. elegans*. Worms were grown for ∼2 days and 5 hours at 20 °C until reaching young adult stage. A summary of the culture methods and drug treatments can be found in S1 Table.

### Device Fabrication

The microfluidic device used in this paper was fabricated by conventional soft lithography technique using polydimethylsiloxane (PDMS). It was reported in our previous paper (S1 Fig)[24]. Briefly, a silicon wafer was used as master mold and patterned using photolithography with the aid of a chrome mask. The mask was developed using a mask writer (Heidelberg Instruments DW66+). Sequel to patterning the silicon wafer, it was etched using deep reactive ion etching thereby creating holes and trenches that served as molds for the micropillars and channels respectively. The difference in heights between the channel and the micropillars was due to the loading effect.

To develop the microfluidic devices, PDMS polymer was prepared by mixing base and curing agents in a 10:1 weight ratio. The mixture was then degassed in a vacuum jar for 10 min, drop-casted over the silicon master mold and cured in an oven for ∼4 hours at 70 °C. The cured PDMS replica was peeled off the silicon mold and carefully bonded on a glass slide after treatment with oxygen plasma for 2 min. Each device had an array of 2 x 12 micropillar array with 17.5 – 23.5 μm in diameter, 36 μm in height and channel depth of 45 μm. S1 Fig summarizes the geometrical details of each device and disease model applied to.

### Image Acquisition and analysis

For the force assay, images and videos of the micropillar deflection due to worm thrashing were captured using an inverted bright-field microscope (Nikon® Eclipse Ti-U) equipped with a CCD camera (Andor® Clara E). The video of each worm was captured for 15 seconds at 100 fps. Video analysis was performed using Kinovea® and deflections were measured every 10 frames (150 data points per micropillar) resulting in a total of 3600 data points per worm. Further details of the experimental set up can be found in our work[24].

The double mutant LS587 strain muscle morphology was examined with phalloidin (ThermoFisher, A12379) to stain actin filaments and evaluated closely under 60X immersion oil objective[24]. For NL5901 and AM725 protein aggregation was assessed through imaging using 3% agarose pads with 10mM levamisole then worms were imaged using 10X magnification with 10ms exposure time[29]. All imaging was carried out using FITC filter under inverted microscope (Nikon® Eclipse Ti-E) equipped with a CCD camera (Andor® iXon Ultra 897 EMCCD).

### Thrashing Force Measurement

The thrashing force of the worms were measured using elastomeric micropillars incorporated within a microfluidic device. The microfluidic device, with the aid of a notch, partially immobilizes a part (head of tail) of the worm while the remainder of the body is allowed to thrash and consequently deflecting the micropillars. The deflection of the micropillars was captured and recorded for 15 secs. Due to the sensitivity of the micropillars, we observed non-linear deflection of same which implied that the conventional Timoshenko beam theory could not be used for the calculation of forces. To calculate the thrashing force from non-linear displacements, we used our custom finite element model (FEM) which we have previously reported[24]. The maximum deflection of each micropillar was measured and the average was taken for the number of worms used. The finite element model was developed using ABAQUS/CAE 2016. The geometry was meshed using 20-noded quadratic hexahedral elements and analysis was performed with ABAQUS/Standard 2016 using a full-Newton direct solver. The effect of the PDMS soft substrate could be neglected, since the study focuses on the relative changes in the thrashing force only. Comparative analysis of the thrashing force was done using two-way ANOVA statistical technique.

### Statistical Analysis

For comparison of thrashing force data, two-way ANOVA with Tukey’s multiple comparison was used. Changes in actin filament morphology of DMD worm samples were analyzed using chi-squared method. Other statistical analyses on the fluorescent image data were performed using student t-test.

## Results

### Two-dimensional analysis protocol

The general workflow of the two-dimensional analysis is shown in Fig 1A. Once cultured on agarose plate and washed off using M9 solution, worms were split into two fractions. One fraction of the worm suspension was used for thrashing force analysis and the other fraction for image analysis. To quantify thrashing force, we used the PDMS-based micropillar force assay device shown in Fig 1B, which used a constriction channel to partially immobilize the worm. This partial immobilization circumvented vision-based tracking and introduced mechanical stimulation, induced by the walls of the constriction channel, on the head of the worm. The remainder of the worm thrashed on the micropillar array and the deflection was captured with a microscope (Fig 1C). Using a custom non-linear finite element model (see details in Methods section) the force exerted by the worms on each micropillar was calculated (Fig 1D). We evaluated the thrashing force exerted by DMD (LS587), ALS (AM725) and Parkinson’s Disease (NL5901) mutant worms compared to wild-type N2. The average thrashing forces were 25.5 ± 0.92 μN (mean ± standard error of the mean, n = 27), 7.9 ± 0.49 μN (n =25), 11.4 ± 1.43 μN (n =25) and 16.4 ± 1.68 μN (n = 25) for wild-type N2, LS587, AM725 and NL5901, respectively (Fig 1E). Compared to the thrashing force of N2, these force values translated to a decrease of 68.7 %, 55.4 % and 35.7 % for LS587, AM725 and NL5901, respectively. As for the second-dimensional analysis, we performed an image analysis using fluorometry to quantify changes in protein aggregation and morphology of the body wall muscles and validated the force measurement data.

**Fig 1:**
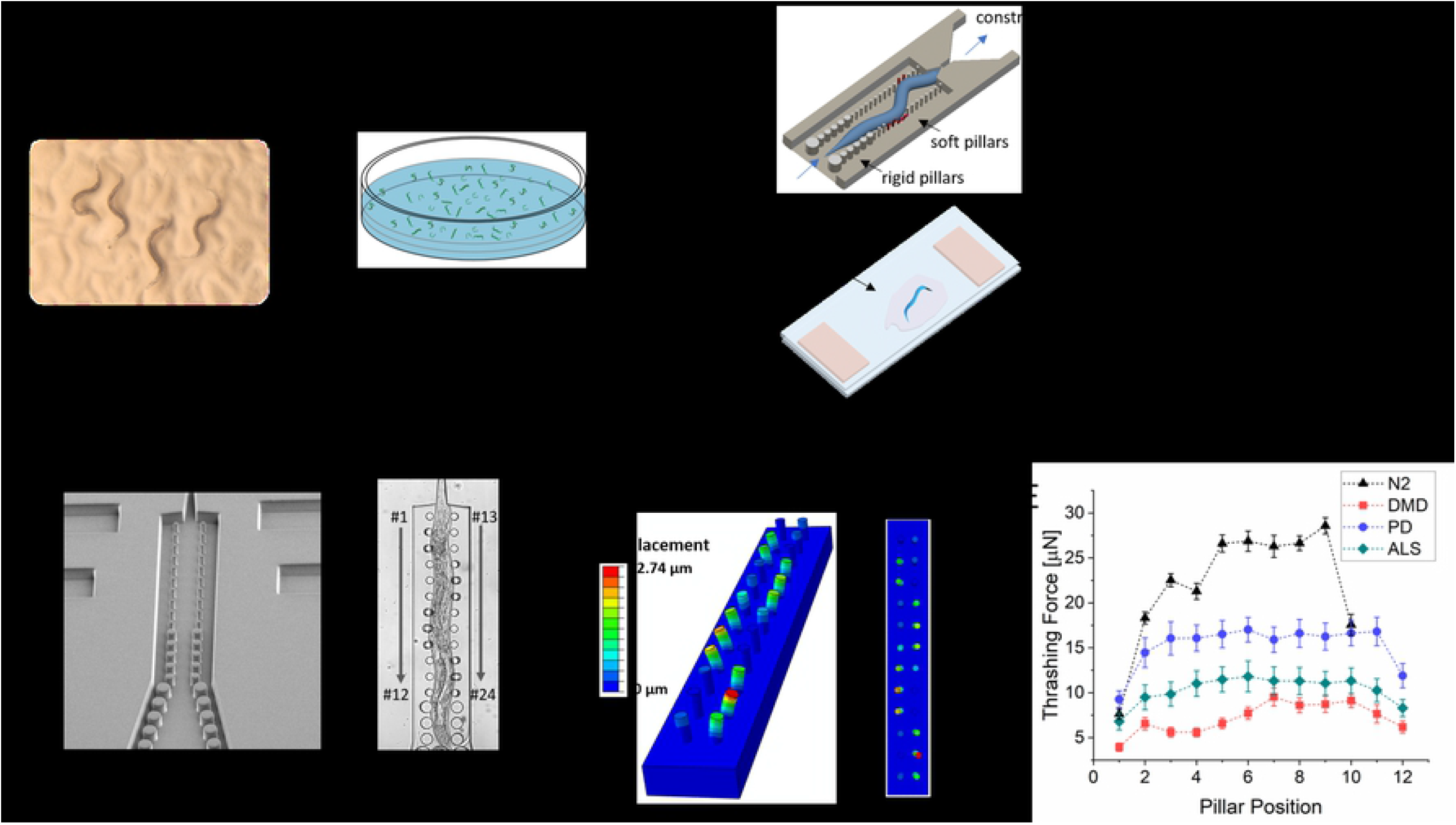
Evaluation of drug efficacy in *C. elegans* using two-dimensional analysis. (A) Protocol for two-dimensional analysis. The first dimensional analysis evaluated the change in thrashing force exerted by the worm before and after drug treatment. The second dimensional analysis used image analysis to evaluate the effect of drug treatment on morphology of the body wall muscles and protein aggregation. (B) Scanning electron micrograph of the force assay chip showing elastomeric micropillars. (C) Optical image showing a partially immobilized young adult worm thrashing on micropillars. (D) Non-linear finite element model used to calculate the thrashing force from the displacements of micropillars. The deflection of micropillars in the FEM image matches that of the optical image in (C). (E) Line graph showing thrashing forces of wild type N2, DMD, ALS and PD model worms in young adult stage. The disease model worms showed a 68.7 %, 55.4 % and 35.7 % decrease in thrashing force for DMD, ALS and PD strains, respectively, compared to the thrashing force of wild-type N2.

### Melatonin and Prednisone improve thrashing force in DMD worm

In *C. elegans*, the *dys-1* gene encodes the protein orthologous to the dystrophin protein associated with DMD in humans which causes progressive muscle loss [30]. To increase the amount of muscle degeneration, *dys-1* mutation has been combined with a hypomorphic mutation, *hlh-1* gene which is a homolog for the *MyoD* gene in human. Using the LS587 (*dys-1;hlh-1)* double mutant worm, we evaluated the effect of two pharmacological treatments, melatonin and prednisone, on the recovery of the thrashing force in our force assay chip. Melatonin has been reported to improve muscle metabolism and strength in mice[31] and clinical trials of DMD patients[31]. Prednisone is the recommended treatment for DMD patients [7] and has been reported to reduce the number of degenerate muscle cells in LS587 mutants[32].

In L4 worms, there was no significant difference in thrashing force with and without treatment for both drugs (Fig 2A and Fig 2B). This insignificant change in thrashing force was expected because L4 worms have only ∼5% degenerate muscle cells[33]. However, in young adult worms, both drugs significantly improved the thrashing force of the worm (Fig 2C and Fig 2D). The average thrashing force around the mid-region of the worm (pillars 5-9) was 9.1 ± 0.94 μN (n = 25) and 9.9 ± 0.35 μN (n = 25) after treatment with melatonin and prednisone, respectively, which translated to a 21.3 % and 40.1 % improvement compared to untreated mutant worms. Proximal to the constriction channel where the head region of the worm was trapped, we observed a 18.8 % and 22.2 % increase in thrashing force for melatonin and prednisone, respectively. These results showed that thrashing force can be used as a phenotype to evaluate drug efficacy in muscle-related disease mutants at both larval and adult developmental stages.

**Fig 2:**
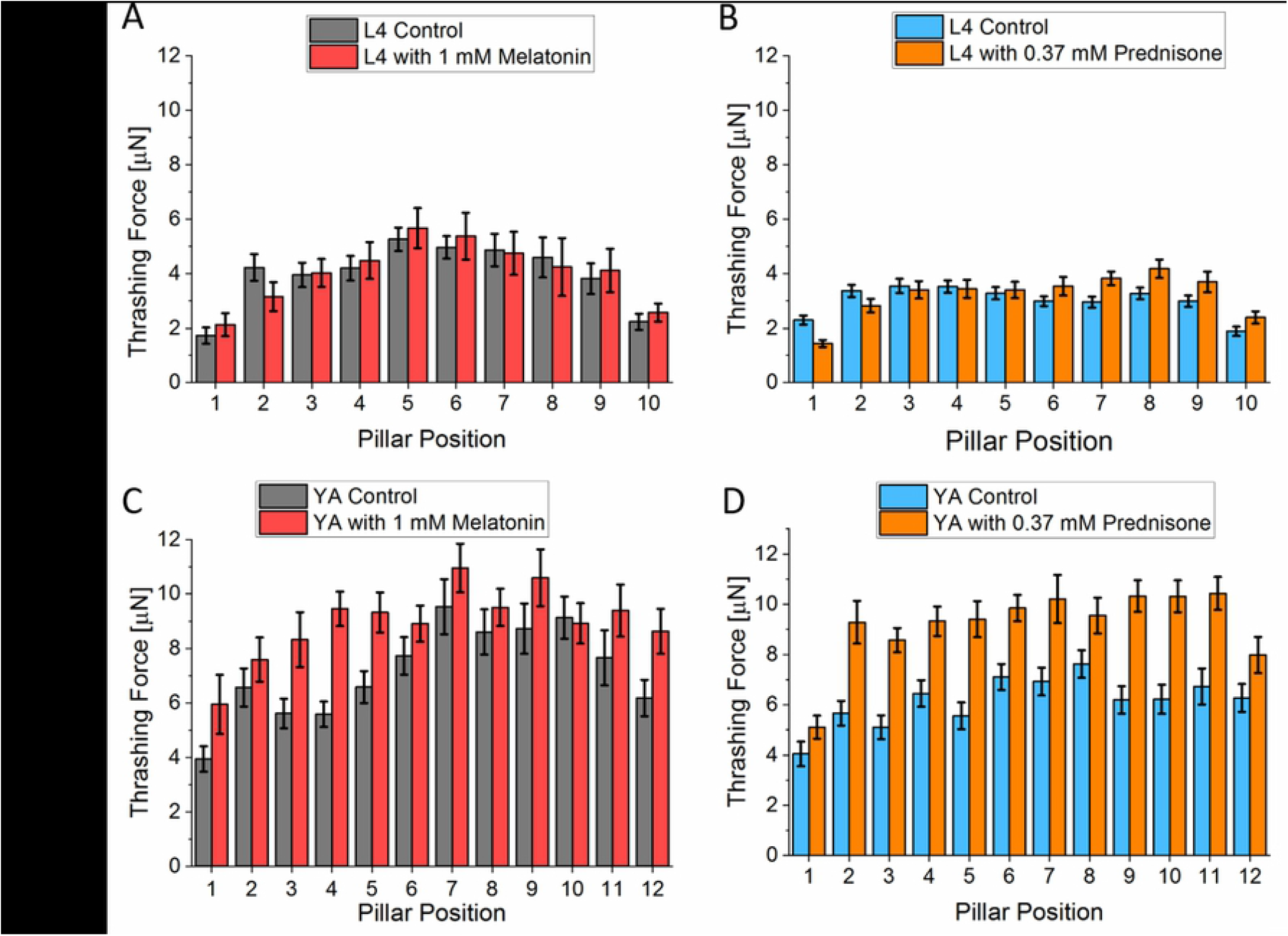
Thrashing force assay in DMD model worm (LS587). (A) L4 stage DMD model worms did not show any significant change in thrashing force when treated with 1 mM melatonin (N=25, p > 0.05). (B) Prednisone treatment on L4 model worms also did not show any change in thrashing force (N=25, p > 0.05). (C) Young adult worms treated with 1 mM melatonin showed significant improvement in thrashing force exerted across all the micropillars when compared to control worms which were not treated (N=25, P < 0.0001). (D) Prednisone treatment also increased the thrashing force of young adult worms compared to worms without any treatment (N=25, p < 0.00001). Significant differences were analyzed using the two-way ANOVA with Tukey’s multiple comparison between average force values of control and treated worms, error bars indicate s.e.m.

To correlate muscle morphology with thrashing force, the sarcomere organization of the body wall muscles was analyzed optically using phalloidin stain[34]. The sarcomere organization was classified using a scoring scale from 0-2 (Fig 3A). A score of 0 indicated healthy parallel and smooth actin filaments while scores 1 and 2 demonstrated wave-like fibers (damaged) with minor waves for score 1 and major damage for score 2[35]. In L4 worms, there was no significant difference in muscle fibers after treatments with both drugs, melatonin (n = 29) and prednisone (n=27) (Fig 3B and Fig 3C). However, there was significant improvement in the morphology of muscle fibers of young adult worms when treated with prednisone (n = 28), with 30 % of worms recovering with score 0 (Figure 3c). Although there was an enhancement in muscle fibers of worms treated with 1 mM melatonin (n=35), it was not a significant difference.

**Fig 3:**
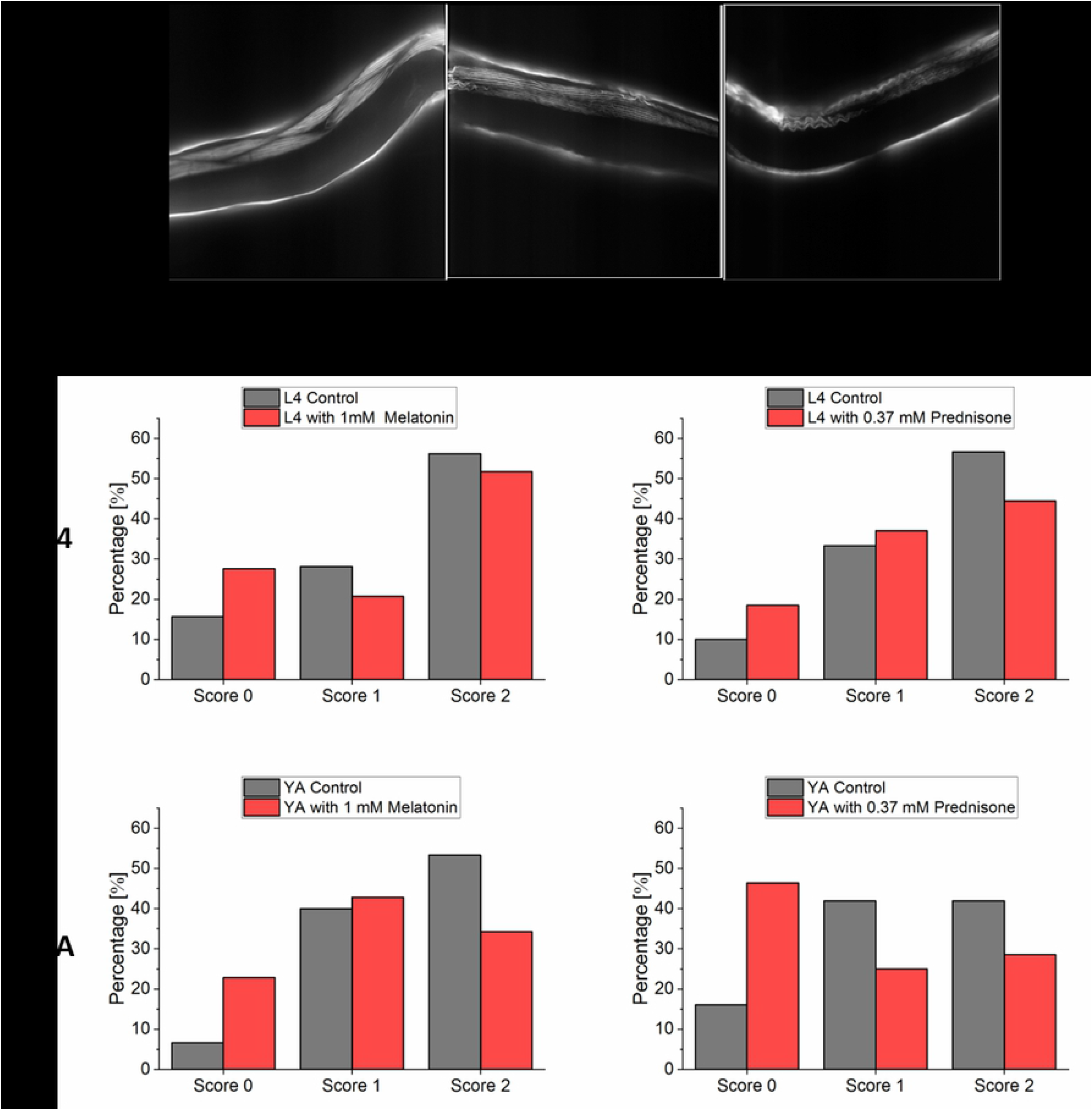
Morphology study of body wall muscle actin filaments for DMD model worm (LS587). The effect of drug treatment with prednisone on muscle morphology in young adult (YA) was studied using phalloidin staining. (A) Score 0 for DMD strain after treatment with prednisone showing smooth and parallel muscle fibers, score 1 for unhealthy muscle with minor waves in control group, and score 2 for damaged actin filaments with major wave like filaments in control group (nontreated). (B) Treatment with 1 mM melatonin for L4 stage and young adult stage both did not show any significant recovery (p = 0.49, p = 0.12). (C) Treatment with 0.37 mM prednisone in L4 stage did not show any significant recovery in muscle morphology (p = 0.54). However, in young adult a significant difference (p = 0.04) was measured. Significant differences were analyzed using chi-squared test. (Melatonin control, L4 stage, N=32, drug treated N=29; Melatonin control, YA, N=30, drug treated N= 35; prednisone control, L4 stage, N=30, drug treated N=27; prednisone control, YA, N=31, drug treated N= 28).

### Riluzole shows dose dependent recovery of thrashing force

As the second disease model, we used a transgenic worm, AM725, for ALS which expresses SOD1 proteins in the body wall muscle cells. We quantified the response of the thrashing force of the worm to two drugs: riluzole and doxycycline. Riluzole has been shown to reduce disease progression and extend patients’ survival by 3 – 6 months[36]. Doxycycline has been used to improve the motility of worms and reduce oxygen consumption[18].

The AM725 mutant worms treated with 30 μM of riluzole did not show any significant change in thrashing force compared to control worms seeded in the absence of the drug. However, when the concentration was increased to 100 μM, there was a significant increase in thrashing force. In the mid region of the worm (micropillars 5 −9), the average of the maximum thrashing force was 11.35 ± 1.5 μN (n = 25) and 14.37 ± 1.44 μN (n = 25) for control worms and those treated with 100 μM riluzole, respectively (Fig 4A), indicating a 26.6 % increase in thrashing force. With doxycycline, we observed an improvement in thrashing force for both concentrations of 10.5 μM and 32 μM. The peak thrashing force, at the mid-region of worm, had average values of 7.13 ± 0.1 μN, 9.06 ± 0.13 μN and 9.38 ± 0.1 μN (n = 27 for all samples) for worms seeded with 0 (control), 10.5 μM and 32 μM doxycycline, respectively (Fig 4B). These results implied a ∼27.1 % and ∼31.6 % increase in thrashing force for 10.5 μM and 32 μM doxycycline treatment, respectively. However, there was no significant difference in the thrashing force between the two doses of doxycycline. Within the reported dosages, only treatment with riluzole was dose-dependent.

**Fig 4:**
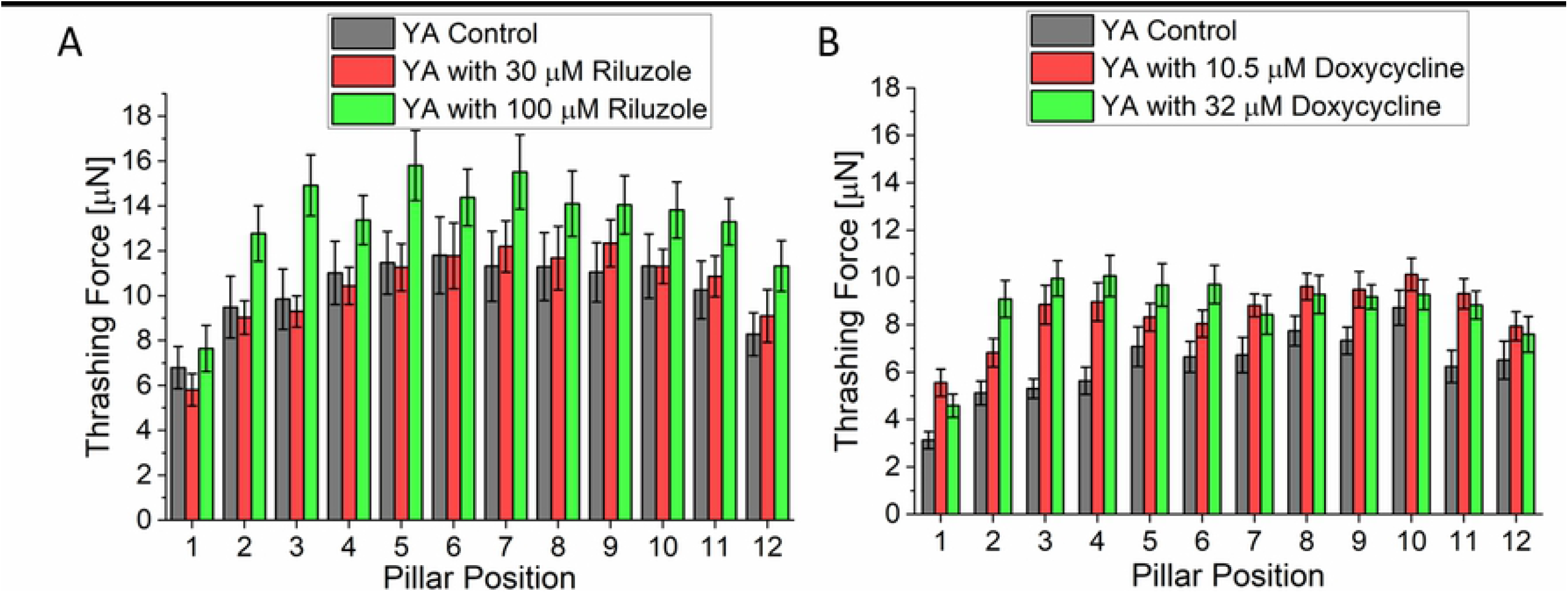
Thrashing force assay in ALS model worm (AM725). (A) Mutant worms treated with 30 µM riluzole did not show any significant change in thrashing force compared to untreated worms (p > 0.05). When the concentration of riluzole was increased to 100 µM, there was a significant increase in the measured thrashing force compared to control worms which were not treated with any drug (N=25, p < 0.001). (B) Treatment with either 10.5 µM or 32 µM of doxycycline significantly improved the thrashing force of AM725 worm compared to control worms (N=27, p < 0.0001). However, there was no significant difference in the thrashing force at the two different concentrations of doxycycline. Significant differences were analyzed using the two-way ANOVA with Tukey’s multiple comparison between average force values of control and treated worms, error bars indicate s.e.m.

The incensement of mutated SOD1 expression in the worm’s body wall muscle cells is an indicator of disease severity[37]. Fig 5A, Fig 5B and Fig 5C show fluorescent images of SOD1 protein aggregate before (n = 34) and after treatment with 30 μM (n = 33) and 100 μM (n=35) riluzole. For the analysis, the size, count, and total area of the protein aggregates were measured (S2 Table). After treatments at both concentrations, there was a significant difference in aggregate size and area. The average size decreased from 69 ± 6.4 to 57 ± 5.4 and 56 ± 4.2, respectively, after treatment with 30 μM and 100 μM riluzole (Fig 5D) while the average area decreased from 0.5 ± 0.04 to 0.39 ± 0.04 and 0.3 ± 0.03, respectively. However, there was no significant difference in aggregates count between control and 30 μM riluzole. However, with 100 μM riluzole, there was a significant difference in average count which dropped from 20 ± 0.9 to 16 ± 0.9. As comparison, the force analysis (Fig 4A) did not show any significant difference between control and 30 μM. In terms of the aggregate average size, there was a decrease for both concentrations from 55 ±2.8 to 50 ± 2.1 and 43 ± 3.1, respectively, but only 32 μM doxycycline decreased significantly (Fig 5E). Doxycycline treatment significantly decreased the average count for treatments with 10.5 μM and 32 μM from 21 ± 0.77 to 19 ± 0.67 and 17 ± 0.65, respectively. For the average area there was no significant decrease for 10.5 μM while treatment with 32 μM doxycycline showed a significant decrease in the average area.

**Fig 5:**
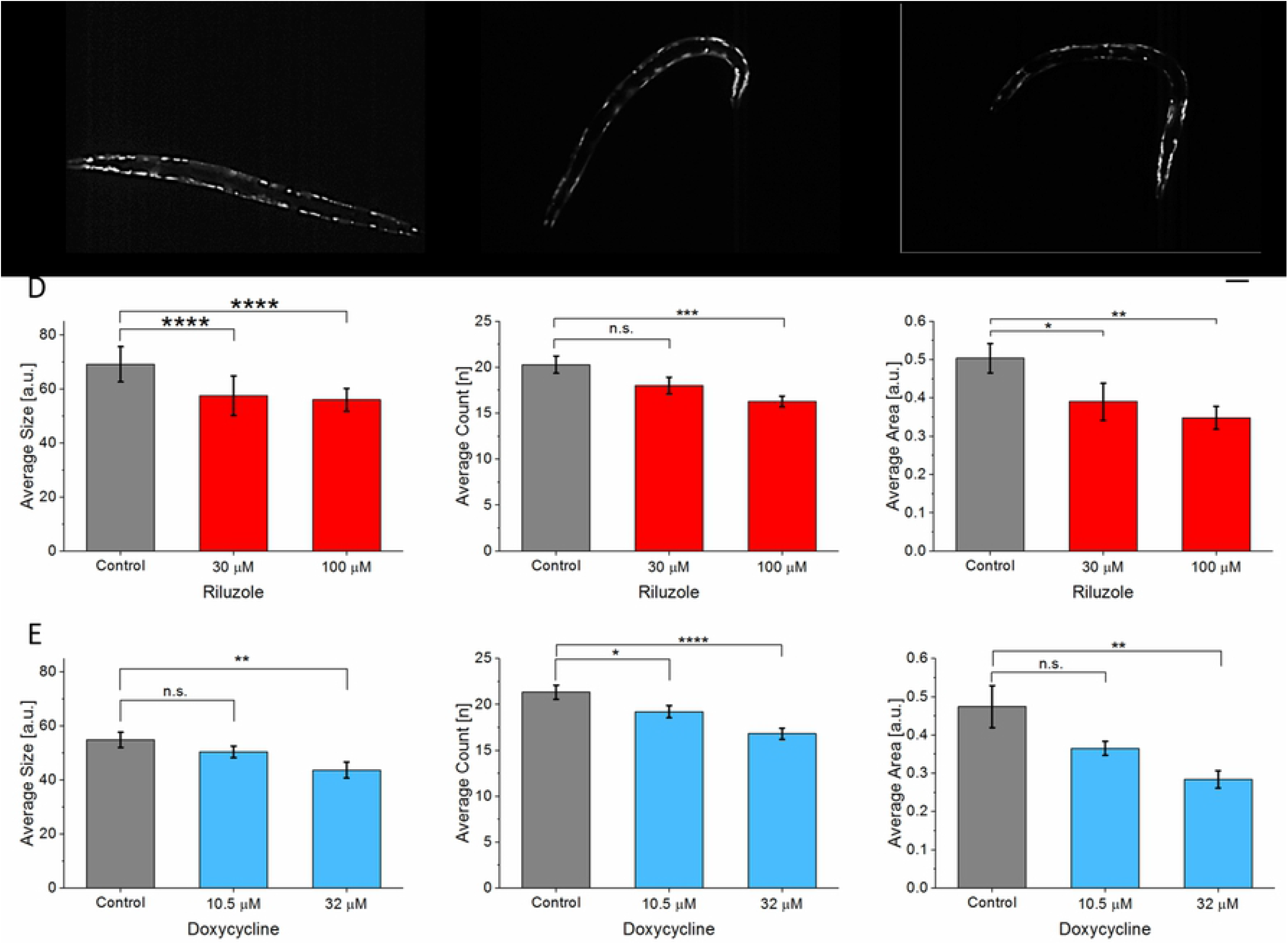
Quantitative analysis of morphology in ALS model (AM725). (A) SOD1 protein aggregates in ALS model before treatment (n = 35). (B) After treatment with 30μM (n = 33). (C) After treatment with 100 μM doxycycline (n = 35). (D) Quantification of protein aggregates after riluzole treatment. 30 μM riluzole showed significant changes in average size and area while its effect on the thrashing force was not significant. (E) After treatment with doxycycline in terms of aggregate size, count, and average area. In the case of doxycycline, 32 μM (n = 40) showed significant changes while 10.5 μM (n = 39) showed no significant changes in terms of average size and area. Significant differences were analyzed using t-test, error bars indicate s.e.m.

### Drug treatment improves thrashing force in PD model worm

Previous study[38] has observed alpha-synuclein (α-synuclein), a small, predominantly presynaptic cytoplasmic protein in the brain of PD patients. Naturally, *C. elegans* does not possess an α-synuclein homolog, however, several transgenic strains have been created such as NL5901 which has human α-synuclein aggregation fused YFP in the muscles of the worm. Using this strain, we quantified the change in thrashing force of the worms after treatment with two pharmacological drugs, pramipexole and levodopa. Pramipexole, a dopamine agonist reported to improve depressive symptoms of PD patients[39], and levodopa, a dopamine precursor which has been the main therapy for PD patients[40].

Treatment of NL5901 worms with either drug showed recovery of thrashing force. Worms treated with 2.5 mM and 5 mM of pramipexole exerted an average thrashing force of 22.09 ± 0.31 μN and 23.15 ± 0.22 μN (Fig 6A) around its mid-region (micropillars 5-9). Compared to the untreated worms (control) which exerted 16.48 ± 0.41 μN (n = 25 for all samples), the results showed 34.1 % and 40.5 % increase in thrashing force for treatments with 2.5 mM and 5 mM of pramipexole, respectively. There was no significant difference in the thrashing force at both drug concentrations. Using levodopa, the average thrashing force increased from 16.34 ± 0.08 μN of the control group to 19.5 ± 0.2 μN and 21.8 ± 0.22 μN (n = 25 for all samples) for worms treated at 0.7 mM and 2 mM, respectively (Fig 6B). This translated to an increment of 19.3 % and 33.3 % with 0.7 mM and 2 mM levodopa treatments, respectively, indicating an increase in thrashing force with higher drug concentration. This result showed a dose-dependent improvement in thrashing force with levodopa while no such dose-dependent recovery was observed with pramipexole.

**Fig 6:**
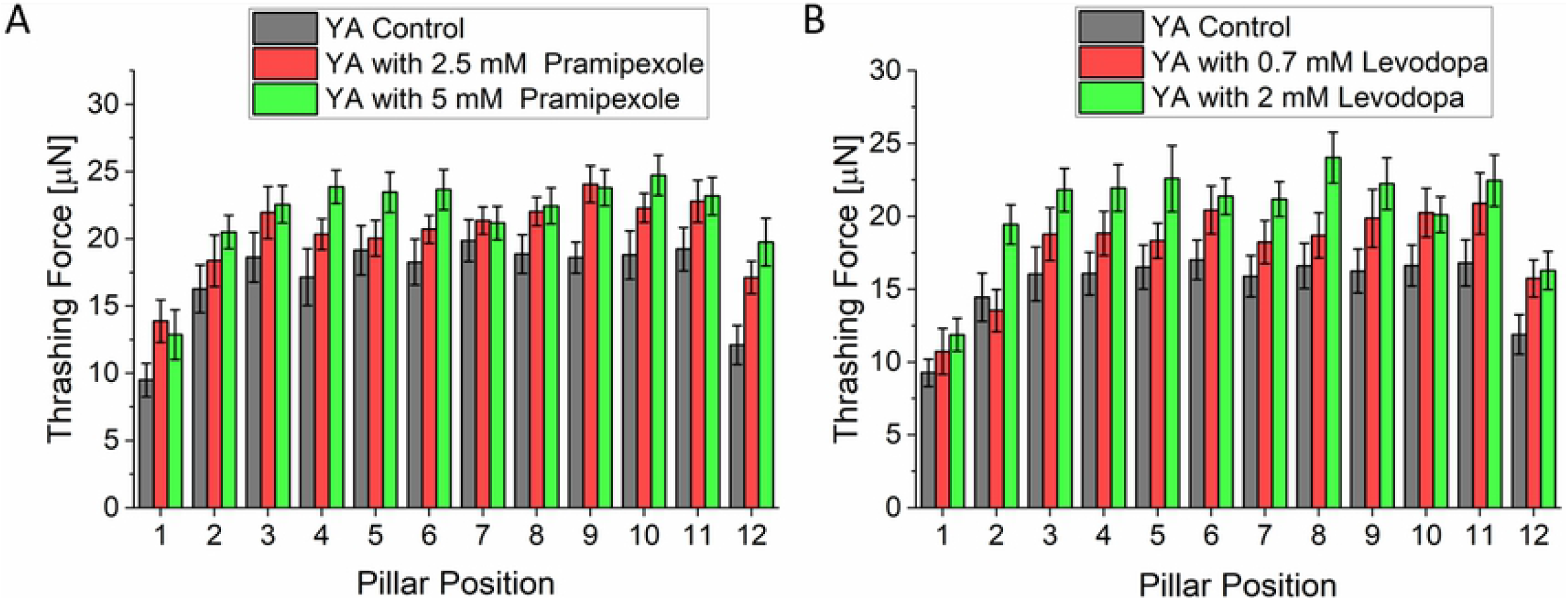
Thrashing force assay in Parkinson Disease model worm (NL5901). (A) Treatment of NL5901 worms with both 2.5 mM or 5 mM of pramipexole significantly improved the thrashing force compared to untreated worms (N=25, p < 0.001). There was no significant difference in thrashing force of worms between the two concentrations of the drug. (B) With levodopa, there was significant change in thrashing force of worms treated with 0.7 mM of the drug. The thrashing force further increased with an increase in levodopa concentration from 0.7 mM to 2 mM (N=25, p < 0.0001). Significant differences were analyzed using the two-way ANOVA with Tukey’s multiple comparison between average force values of control and treated worms, error bars indicate s.e.m.

In NL5901 *C. elegans* strain, the levels of α-synuclein was measured by estimating the fluorescence intensity of YFP in the muscle cells[27]. When the strain was treated with pramipexole, there no significant difference in fluorescence intensity between control and 2.5 mM treated worms (n = 27) (Fig 7D). However, treatment with 5 mM pramipexole (n = 24), there was significant decrease in the fluorescence intensity of the protein aggregates. Treatment with 0.7mM (n = 32) and 2mM (n = 31) levodopa did not show any significant difference in fluorescence signal intensity (Fig 7E).

**Fig 7:**
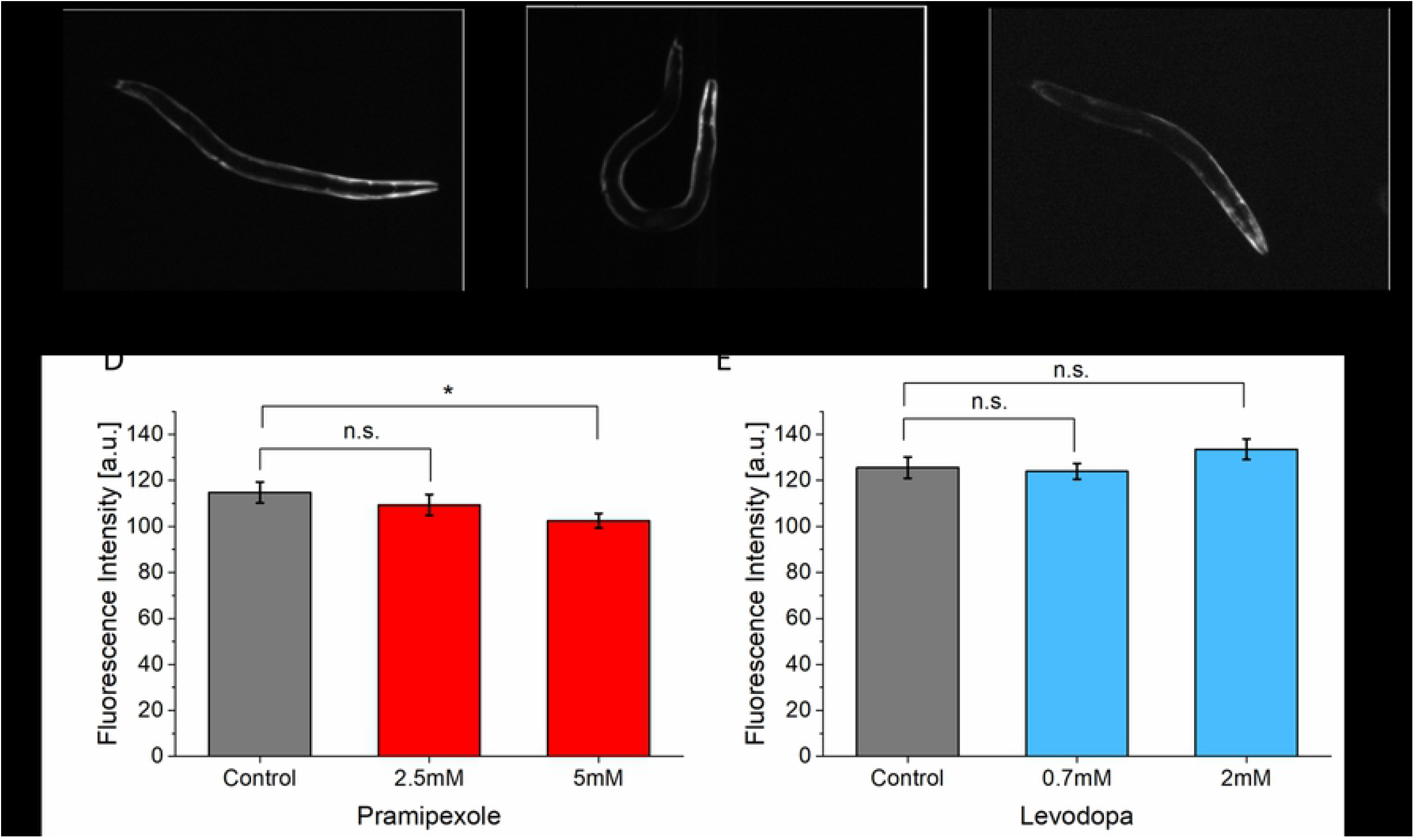
Quantification of fluorescence intensity of α-synuclein protein in NL5901. (A) Fluorescent image of α-synuclein protein accumulation before treatment (n = 31). (B) after treatment with 2.5 mM (n = 27). (C) after treatment with 5 mM (n = 24) pramipexole. (D) Fluorescence intensity of protein accumulation before and after treatment with pramipexole. (E) Fluorescence intensity of protein accumulation before (n = 35) and after treatment with levodopa 0.7mM (n=32) and 2mM (n=31). Significant differences were analyzed using t-test, error bars indicate s.e.m.

## Discussion

### Duchenne Muscular Dystrophy

In this study, we treated *C. elegans* DMD mutant (LS587) with prednisone and melatonin, and observed significant improvement in the thrashing force of young adult worms. However, there was no change observed in L4 worms as expected. A possible explanation for this result could be due to non-degeneration of muscle cells until after L4 stage as a result of cellular repairs which occur at the end of each larval stage and delay muscle cell damages till adulthood[32]. The significant improvement in young adult worms aligned with the prior studies that treated single mutant *dys-1*[14] and double mutant *dys-1;hlh-1*[32] worms with prednisone and/or melatonin. In particular, our results with partially immobilized worms were in close agreement with the outcomes of the study conducted Hewitt et. al.[14]which used the same dosage of drug treatment on free-moving *dys-1(eg33)* mutant worms. Patients treated with melatonin have been reported to decrease serum creatine kinase which indicates reduced muscle damage and oxidative stress[31]. Similar reduction in oxidative stress and improved redox status have also been reported in mdx mice treated with melatonin[31]. Prednisone is known to act against inflammatory processes in the muscles of DMD patients[41], however this cannot be investigated in *C. elegans* due to the lack of inflammatory-mediated amplification in the worm’s degenerative muscles. Another and a study suggested the sarcolemma stabilization due to little alterations in plasma membrane of muscle cells as another mechanism through which prednisone acts[32]. The image analysis of muscle morphology for DMD model supported the thrashing force data that there was no significant difference observed until L4 stage for both drugs, melatonin and prednisone. Even worms treated with melatonin until young adult stage did not show a significant recovery in muscle morphology while the thrashing force assay detected a significant increase of the thrashing force once treated. In the case of prednisone, however, its muscle morphology improved significantly when treated until young adult stage. It has been reported that LS587 mutant treated with prednisone recovered its muscle morphology almost to healthy state[33]. This result suggested that the thrashing force analysis was more sensitive than the image analysis when detecting the efficacy of drugs.

### Amyotrophic Lateral Sclerosis

Both riluzole, an internationally approved ALS drug, and doxycycline, an antibiotic, were able to improve the thrashing force of *C. elegans*. The specific mechanism through which riluzole acts is still unknown, but several neuroprotective properties have been ascribed to it, such as inhibition of presynaptic glutamate release and upregulation of the expression of growth factors[42,43]. Doxycycline has been reported to improve locomotion in worms through activation of mitochondrial unfolded protein (UPR^mt^)[18]. A potential explanation for this working mechanism of doxycycline is due to the fact that mitochondrial accumulation of misfolded *SOD1* has been reported as a potential trigger for motor neuron death. Our result was in agreement with the previous studies that have shown amelioration of loss of motility in ALS mutant worms through treatment with riluzole[13] and doxycycline[18]. The dose dependence of the thrashing force of worms treated with riluzole at 30 μM and 100 μM corroborated with the results from the previously reported study on worm speed[13]. Doxycycline significantly improved the thrashing force at both concentrations, 10.5 μM and 32 μM. When evaluating the protein aggregates in terms of count, size and average area, riluzole, was more effective with higher dosage of 100 μM compared to 30 μM in agreement with the thrashing force measurement. The average count of aggregates decreased significantly after treatment with 100 μM of riluzole but not with 30 μM. In terms of average size and area, there was a significant decrease when treated with both 30 μM and 100 μM of riluzole. In comparison, the force measurement showed no effect of the drug at 30 μM. This result implied that the image analysis can also be more sensitive to the changes due to drug treatment than the force analysis. This finding proved the complementarity of both analyses when quantifying the drug efficacy. Doxycycline has been reported to decrease the size of *SOD1* protein aggregate but not the aggregate count for SJ4100 strain[18]. In this study, doxycycline treatment decreased both the size and count of the protein aggregates significantly at 32 μM but not at 10.5 μM whereas the force analysis could discern a subtle change at the lower drug concentration.

### Parkinson’s Disease

Drug treatment of the PD model with pramipexole and levodopa showed an improvement in thrashing force for both drugs. However, pramipexole showed a better recovery in thrashing force compared to levodopa. In clinical trials, pramipexole has been reported to slow down the onset of dopaminergic neurons in PD patients thereby improving the symptoms of the disease[39]. Levodopa is the leading treatment for PD and has proved highly effective in ameliorating symptoms of the disease[40]. The improvement of thrashing force exerted by the worms treated with levodopa agrees with the prior studies on the locomotory parameters of PD model worms with similar treatment[44]. A potential explanation for the effect of levodopa is the increase in polarized distribution and expression of type-1 dopamine receptors in acetylcholinergic motor neurons[44] compared to untreated worms. Our result showed that the improvement of thrashing force is not dependent on drug dosage for pramipexole at the concentration levels used in this study. For levodopa, however, there was a dependency showing higher thrashing force at 2 mM compared to 0.7 mM. To the best of our knowledge, this was the first time that the effect of pramipexole has been studied on a locomotory parameter of PD *C. elegans* model. This result underscored the potential of pramipexole as treatment for PD considering the complications due to chronic use of levodopa[40]. When analyzing the protein accumulation by measuring the fluorescence intensity of YFP in the muscle cells, the fluorescence intensity decreased when treated with 5 mM pramipexole, but no significant difference was observed when treated with 2.5 mM. When treated with both concentrations of levodopa, 0.7 mM and 2 mM, there was no significant difference measured. Compared to the thrashing force analysis which showed significant improvement with treatment of pramipexole or levodopa at both concentrations, the image analysis seemed to be less sensitive. In all three disease model cases, the force analysis seemed to be generally more sensitive than the image analysis and delivered a quantitative readout less ambiguous compared to the analysis of fluorescent images. However, our study also confirmed that both the thrashing force as well as the image analysis should be conducted together to validate the measurement data and to detect subtle changes in complementarity since one of them might be more sensitive than the other one depending on the disease models such as in ALS model.

## Conclusion

To evaluate the efficacy of drug treatment on three *C. elegans* models for neuromuscular diseases, we have implemented a two-dimensional workflow analysis consisting of a thrashing force measurement in a microfluidic chip and an image analysis using an agarose pad. In the first-dimensional analysis, we evaluated thrashing force of these disease model worms before and after drug treatment, while in the second-dimensional analysis, we performed a quantitative image analysis of the protein aggregation and morphological studies of the body wall muscles. The thrashing force analysis was more sensitive to measure the changes resulting from drug treatment compared to the image analysis as demonstrated in the case of DMD and PD models and partially in ALS model. To the best of our knowledge, this was the first study that reported the force exerted by the body wall muscles of ALS and PD *C. elegans* models. All these results underlined the potential of our force assay chip in screening of potential drug candidates for the treatment of DMD, ALS, PD and potentially other muscle-related diseases. Our partial immobilization-based device reduced the complexities of instrumentation associated with tracking worms through computer vision. which offers scalability for multiplexing by using multiple parallel channels in the force assay chip. This multiplexing step in combination with an imaging chip for *C. elegans* could ultimately lead to higher throughput for drug screening.

## Acknowledgements

The authors will like to acknowledge support from Dr. Hala Fahs, Suma Gopinathan and Fathima Refai of the Center for Genomics and Systems Biology, NYU Abu Dhabi in sourcing the *C. elegans* strains used in this work. We acknowledge Navajit Baban’s support in drawing the 3D schematic images. We are also thankful for the support by NYUAD microfabrication core facility for the device fabrication. This work was supported by the Al Jalila Foundation [AJF201633]. S. Sofela was supported by the NYUAD Global PhD Fellowship program.

## Supporting Information

**S1 Fig, The microfluidic chip for quantifying thrashing force exerted by *C. elegans***. (A) Optical image showing two developmental stages of worms (L4 and young adult) thrashing on the PDMS-based micropillars. The deflection of the micropillars was used to quantify the thrashing force exerted by the worm. (B) Table showing geometric parameters for force assay chip and diameters of worms used in this study.

**S1 Table, Summary of worm culture and drug treatments**.

**S2 Table, Summary of significance test of SOD1 protein aggregate quantification before and after drug treatment**.

